# Evolutionary forces affecting synonymous variations in plant genomes

**DOI:** 10.1101/086231

**Authors:** Yves Clément, Gautier Sarah, Yan Holtz, Felix Homa, Stéphanie Pointet, Sandy Contreras, Benoit Nabholz, François Sabot, Laure Sauné, Morgane Ardisson, Roberto Bacilieri, Guillaume Besnard, Angélique Berger, Céline Cardi, Fabien De Bellis, Olivier Fouet, Cyril Jourda, Bouchaib Khadari, Claire Lanaud, Thierry Leroy, David Pot, Christopher Sauvage, Nora Scarcelli, James Tregear, Yves Vigouroux, Nabila Yahiaoui, Manuel Ruiz, Sylvain Santoni, Jean-Pierre Labouisse, Jean-Louis Pham, Jacques David, Sylvain Glémin

## Abstract

Base composition is highly variable among and within plant genomes, especially at third codon positions, ranging from GC-poor and homogeneous species to GC-rich and highly heterogeneous ones (particularly Monocots). Consequently, synonymous codon usage is biased in most species, even when base composition is relatively homogeneous. The causes of these variations are still under debate, with three main forces being possibly involved: mutational bias, selection and GC-biased gene conversion (gBGC). So far, both selection and gBGC have been detected in some species but how their relative strength varies among and within species remains unclear. Population genetics approaches allow to jointly estimating the intensity of selection, gBGC and mutational bias. We extended a recently developed method and applied it to a large population genomic datasets based on transcriptome sequencing of 11 angiosperm species spread across the phylogeny. We found that base composition is far from mutation-drift equilibrium in most genomes and that gBGC is a widespread and stronger process than selection. gBGC could strongly contribute to base composition variation among plant species, implying that it should be taken into account in plant genome analyses, especially for GC-rich ones.

## Introduction

Base composition strongly varies across and within plant genomes [1]. This is especially striking at the coding sequence level for synonymous sites where highly contrasted patterns are observed. Most Gymnosperms, basal Angiosperms and Eudicots have relatively GC-poor and homogeneous genomes. In contrast, Monocot species present a much wider range of variation from GC-poor species to GC-rich and highly heterogeneous ones, some with bimodal GC content distribution among genes, these differences being mainly driven by GC content at third codon position (GC3) [1]. Commelinids (a group containing palm trees, banana and grasses, among others) have particularly GC-rich and heterogeneous genomes but GC-richness and bimodality have been showed to be ancestral to Monocots, suggesting erosion of GC content in some lineages and maintenance in others [2]. As a consequence, in most species, synonymous codons are not used in equal frequency with some codons more frequently used than others, a feature that is called the codon usage bias [reviewed in 3]. This is true even in relatively homogeneous genomes such as in *Arabidopsis thaliana* [e.g. 4].

Which forces drive the evolution of genome base composition and codon usage is still under debate. Mutational processes can contribute to observed variations between species and within genomes [e.g. 5]. However, mutation can hardly explain a strong bias towards G and C bases, as it is biased towards A and T in most organisms studied so far [Chapter 6 in 6]. Selection on codon usage (SCU) has thus appeared as one of the key forces shaping codon usage as it has been demonstrated in many organisms both in prokaryotes and eukaryotes [reviewed in 3]. Codon bias can thus result from the balance between mutation, natural selection and genetic drift [7]. The main cause for SCU is likely that preferred codons increase the accuracy and/or the efficiency of translation but other mechanisms involving mRNA stability, protein folding, splicing regulation and robustness to translational errors could also play a role [3,8,9]. In some species, SCU appears to be very weak or inexistent, typically when effective sizes are small [10], as typically assumed for mammals [but see 8]. However, mammalian genomes exhibit strong variations in base composition, the so-called isochore structure [11], which are mainly driven by GC-biased gene conversion (gBGC) [12]. gBGC is a neutral recombination-associated process favouring the fixation of G and C (hereafter S for strong) over A and T (hereafter W for weak) alleles because of biased mismatch repair following heteroduplex formation during meiosis [13]. Although gBGC is a neutral process – *i.e.* the fate of S vs W alleles is not driven by their effect on fitness – gBGC induces a transmission dynamic during reproduction identical to natural selection for population genetics [14]. Therefore, we here refer to it as a “selective-like” process as opposed to mutation and drift. gBGC has been experimentally demonstrated in yeast [15,16], humans [17,18], birds [19] and rice [20]. Many indirect genomic evidences also supported gBGC in eukaryotes [21,22] and even recently in some prokaryotes [23].

In plants, both SCU [4,24,25] and gBGC [21,26,27] have been documented, but how their magnitudes and relative strength vary among species remains unclear. Recently, it has been proposed that the wide variations in genic GC content distribution observed in Angiosperms could be explained by the interaction between gene structure, recombination pattern and gBGC [28]. Increasing evidence suggests that in various organism, including plants, recombination occurs preferentially in promoter regions of genes, or near transcription initiation sites [29,30,31], generating a 5’-3’ recombination gradient, and consequently a gBGC gradient. A mechanistic consequence is that short genes, especially with no or few introns, are on average GC-richer [32]. A stronger gBGC gradient and/or a higher proportion of short genes would increase the average GC content and simple changes in the gBGC gradient can explain a wide range of GC content distribution from unimodal to bimodal ones [28].

So far, the magnitude of gBGC and SCU has only been quantified in a handful of plant species [24,25,27,33]. As in other species studied, weak SCU and gBGC intensities were estimated. The population-scale coefficients, 4*N*_*e*_*s* or 4*N*_*e*_*b*, are usually of the order of 1, where *N*_*e*_ is the effective population size and *s* and *b* the intensity of SCU and gBGC respectively [24,25,27,33,34,35]. However, high gBGC values (4*N*_*e*_*b* > 10) have been estimated in the close vicinity of recombination hotspots in mammals [33,36] and across the entire genome in honeybee [37]. Differences in population-scale intensities can be due to variation in *N*_*e*_ and/or in *s* or *b*. For gBGC, *b* is the product of the recombination rate *r* and the basal conversion rate per recombination event, *b*_0_. Within a genome, variations in gBGC intensities are mainly due to variation in recombination rate [e.g. 33]. Among species, *b*_0_ can also vary. For instance, *b* was estimated to be 2.5 times lower in honeybees than in humans but recombination rate is more than 18 times higher [37], suggesting that *b*_0_ could be 45 times lower in honeybees than in humans. The very intense population-scale gBGC in honeybees is thus explained by the combination of a large *N*_*e*_ and extremely high recombination rates [37].

Several methods have been developed to estimate the intensity of SCU and gBGC, either from polymorphism data alone, or from the combination of polymorphism and divergence data [e.g. 33,35,38]. These methods rely on the fact that preferred codons (for SCU) or GC alleles (for gBGC) are expected to segregate with higher frequency than neutral and un-preferred or AT alleles, fitting a population genetics model with selection or gBGC to the different site frequency spectra (SFS). As demography affects SFS, it must be taken into account in the model. Moreover, mutations must be polarized, *i.e.* the ancestral or derived state of mutations must be determined using one or several outgroup species. Otherwise, selection or gBGC can be estimated from the shape of the folded SFS only under the assumption of equilibrium base composition [e.g. 35,38], which is not the case in mammals [39] and some Monocots [2], for example. As errors in the polarization of mutations can lead to spurious signatures of selection or gBGC [40], this issue must also be taken into account.

Here we used and extended the recent method developed by Glémin et al. [33] that controls for both demography and polarization errors. We applied it to a large population genomic dataset of 11 species spread across the Angiosperm phylogeny to detect and quantify the forces affecting synonymous positions. We specifically address the following questions: (i) is base composition mainly affected by neutral or selective forces? (ii) if active, what are the intensities of gBGC and SCU and how do they vary across species? (iii) are the average gBGC and the 5’-3’ gBGC gradient stronger in GC-rich genomes? Our results show that base composition is far from mutation-drift equilibrium in most genomes, that gBGC is a widespread process being the major force acting on synonymous sites, overwhelming the effect of SCU and contributing to explain the difference between GC-rich (Commelinids, here) and GC-poor genomes (Eudicots and yam, here).

## Results

### Building a large dataset of sequence polymorphism and divergence in 11 plant species

We focused on 11 plant species spread across the Angiosperm phylogeny with contrasted base composition and mating systems (Figure 1 and Table 1). To survey the wide variation observed in Monocots, and in line with the sampling of a previous study [2], we sampled one basal Monocots (*Dioscorea abyssinica*, yam), two non-grass Commelinids (*Musa acuminata*, banana and *Elaeis guineensis*, palm tree) and three grasses with contrasted mating system (*Pennisetum glaucum*, pearl millet, *Sorghum bicolor*, sorghum and *Triticum monococcum*, einkorn wheat). In Eudicots, both Rosids (*Theobroma cacao*, cacao and *Vitis vinifera*, grapevine) and Asterids (*Coffea canephora*, coffee tree, *Olea europaea*, olive tree and *Solanum pimpinellifolium*, tomato) are represented. For practical reasons cultivated species have been chosen but we only sampled wild individuals over the species range, except for palm tree for which cultivated individuals were sampled (See Table S1 for sampling details). In this species cultivation is very recent without real domestication process (19th century [41]). For each species we used RNA-seq techniques to sequence the transcriptome of about ten individuals plus two individuals from two outgroup species, giving a total of 130 individual transcriptomes. When a well annotated reference genome was available (see Material and Methods), we used it as a reference to map sequenced reads, otherwise we used a *de novo* transcriptome assembly already obtained for these species (focal + outgroups) [42] (Table. 1 and S2). After quality trimming and mapping of the raw reads, we kept contigs with at least one read mapped in every individual. This initial dataset was used for gene expression analyses (see below). Genotype calling and filtering of paralogous sequences were performed using the *read2snp* software [43] for each species separately, and coding sequence regions were extracted (see Material and Methods). The resulting datasets were used to compute diversity statistics that did not require any outgroup information. The number of identified SNPs varies from 4,409 in *T. monococcum* (which suffered from the lowest depth of sequencing) to 115,483 in *C. canephora*. Variations in the numbers of SNPs also revealed the large variation in polymorphism levels with πS ranging from 0.17% in *E. guineensis* to 1.22% in *M. acuminata*. The level of constraints on proteins, as measured by the π_N_/π_S_ ratio, varies between 0.122 in *T. monococcum* and 0.261 in *E. guineensis* (Table 2). For the analyses requiring polarized SNPs, we also added orthologous sequences from two outgroups to each sequence alignment of the focal species individuals (see Material and Methods). The number of polarized SNPs ranged from 3,253 in *S. pimpinellifolium* to 89,793 in *M. acuminata*. Other details about the datasets are given in Table 2. Overall, the dataset does not represent the full transcriptome of each species but allows large-scale comparative analyses.

**Figure 1:**
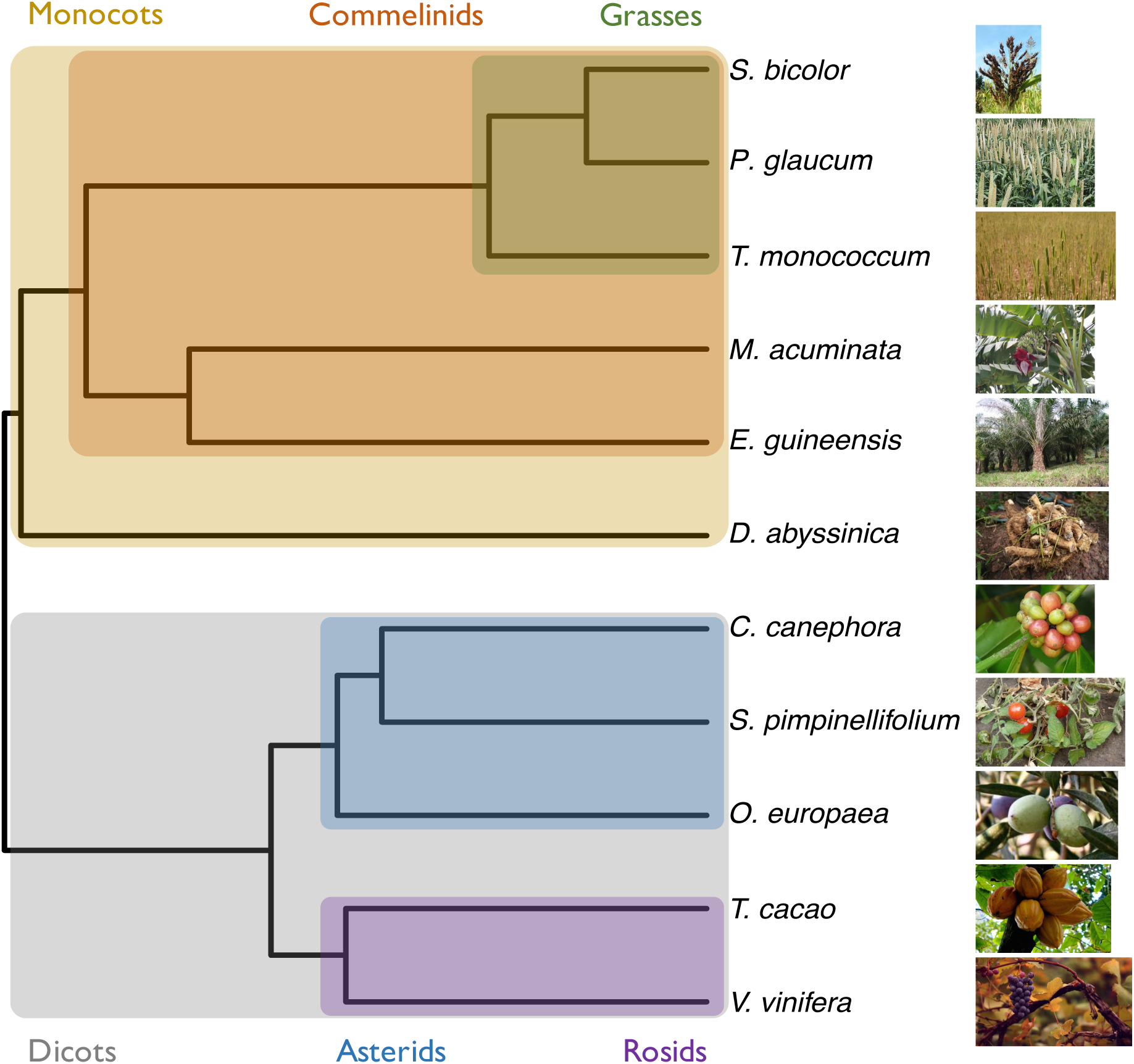
Phylogeny of the species used in this study. Phylogenetic relationship of the species in this study. The phylogeny was computed with PhyML [69] on a set of 33 1S1 orthologous protein clusters obtained with SiLiX [70]. All images were obtained from the wikicommons website.

**Table 1:** List of studied species and datasets characteristics.

| Species | Name | Group | Mating system | Outgroup 1 | Outgroup 2 | Reference | # of individuals |
| --- | --- | --- | --- | --- | --- | --- | --- |
| *Sorghum bicolor* | Sorghum | Monocot - Commelinid | Mixed | *Sorghum brachypodium* | *Zea mays* | Genome | 9 |
| *Pennisetum glaucum* | Pearl millet | Monocot - Commelinid | Outcrossing | *Pennisetum polystachion* | *Pennisetum alopecuroides* | Transcriptome | 10 |
| *Triticum monococcum* | Einkorn wheat | Monocot - Commelinid | Selfing | *Taeniatherum caput-medusae* | *Eremopyrum bonaepartis* | Transcriptome | 10 |
| *Musa acuminata* | Banana | Monocot - Commelinid | Outcrossing | *Musa balbisiana* | *Musa becarii* | Transcriptome | 10 |
| *Elaeis guineensis* | Oil palm tree | Monocot - Commelinid | Outcrossing | *Phoenix dactylifera* | *Mauritia flexuosa* | Transcriptome | 10 |
| *Dioscorea abyssinica* | Yam | Monocot - Basal | Outcrossing | *Dioscorea praehensilis* | *Dioscorea trifida* | Transcriptome | 5 |
| *Coffea canephora* | Coffee tree | Eudicot - Asterid | Outcrossing | *Empogona ruandensis* | *Coffea pseudozanguebariae* | Transcriptome | 12 |
| *Solanum pimpinellifolium* | Tomato | Eudicot - Asterid | Mixed | *Solanum melongena* | *Capsicum annuum* | Genome | 10 |
| *Olea europaea subsp. europaea* * | Olive tree | Eudicot - Asterid | Outcrossing | *Olea europaea subsp. cuspidata* | *Phillyrea angustifolia* | Transcriptome | 10 |
| *Theobroma cacao* | Cocoa | Eudicot - Rosid | Outcrossing | *Herrania nitida* | *Theobroma speciosa* | Genome | 10 |
| *Vitis vinifera* | Grape vine | Eudicot - Rosid | Outcrossing | *Vitis romaneti* | *Vitis riparia* | Genome | 12 |

**Table 2:**
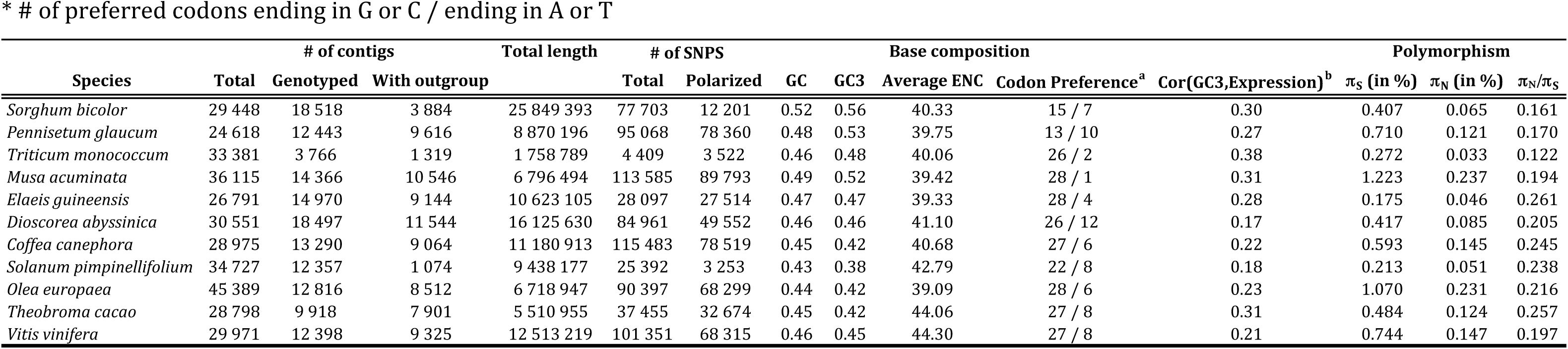
Global statistics for each dataset. GC and GC3 have been computed on the total number of contigs

### Base composition, patterns of codon usage and codon preferences vary across species

We first looked at base composition: GC3 varies from 0.38 to 0.44 in Eudicots and from 0.46 to 0.56 in Monocots (Table 2). As observed in previous studies [2,39], these values tend to be lower than genome wide averages (when available) but the relative differences in base composition among species are conserved, notably the GC-poorness of Eudicots compared to Monocots. Grass species exhibit a bimodal GC3 distribution except *T. monococcum* where bimodality is not apparent (Figure S1). This is likely because the sequencing depth was lower for this species so that GC-rich genes (most likely short ones) have been under sampled. We also characterized codon usage in each species by computing the Relative Synonymous Codon Usage (RSCU) for every codon as the frequency of a particular codon normalised by the frequency of the amino-acid it codes for (Table S3, Figure S2). Patterns of RSCU are relatively consistent between species but reflect differences of GC content between them, notably a higher usage of G or C-ending codons in GC-rich species.

In order to evaluate the possible effect of selection on codon usage, we defined the sets of preferred (P) and un-preferred (U) codons for each species. The fitness consequences of using optimal or suboptimal codons should be higher in highly expressed genes, causing the usage of optimal codons to increase with gene expression (and that of non-optimal ones to decrease). Thus, we defined preferred (resp. un-preferred) codons as codons for which RSCU increases (resp. decreases) with gene expression as in [44] (see Materials & Methods for more details). Table S3 shows detailed results for each species. In contrast with genome-wide codon usage, nearly all species show a bias towards preferred codons ending in G or C (Table 2, Figure 2 and Table S3), only *P. glaucum* and *S. bicolor* showing a more balanced AT/GC sharing of codon preference. Preferences for two-fold degenerated codons are highly conserved across species, with only GC-ending preferred codon except for aspartic acid and tyrosine in *P. glaucum* (Figure 2, Table S3). Preferences for other amino-acids are slightly more labile but there are always one preferred GC-ending and one un-preferred AT-ending codon common to all species. Frequency of optimal codons of a gene (Fop, *i.e.* the frequency of preferred codons [45]), increases with expression as expected but the difference in Fop between the most highly and most lowly expressed genes is weak to moderate (from ~5% in *C. canephora* to 15% in *T. monococcum* and *M. acuminata*) and tends to be higher in Commelinid species (Figure 3). Because most preferred codons end with G or C, GC3 and expression are also positively correlated in all species.

**Figure 2:**
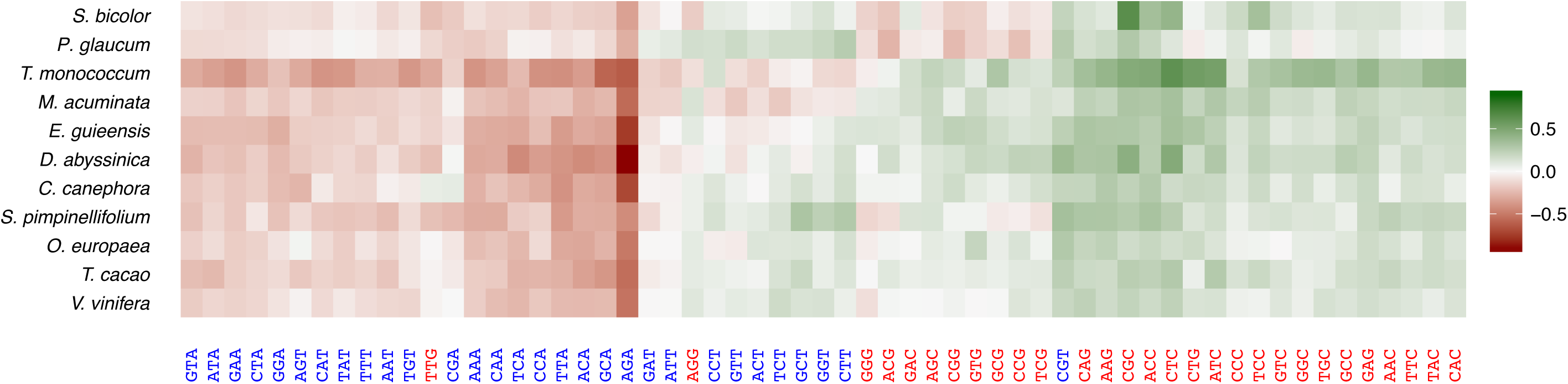
Patterns of codon preference among the 11 studied species. The colour scale indicates the magnitude of ΔRSCU, the difference in the Relative Synonymous Codon Usage between highly and lowly expressed genes. The greenest codons are the most preferred and the reddest the least preferred. Codons ending in G or C are in red and those ending in A or T in blue.

**Figure 3:**
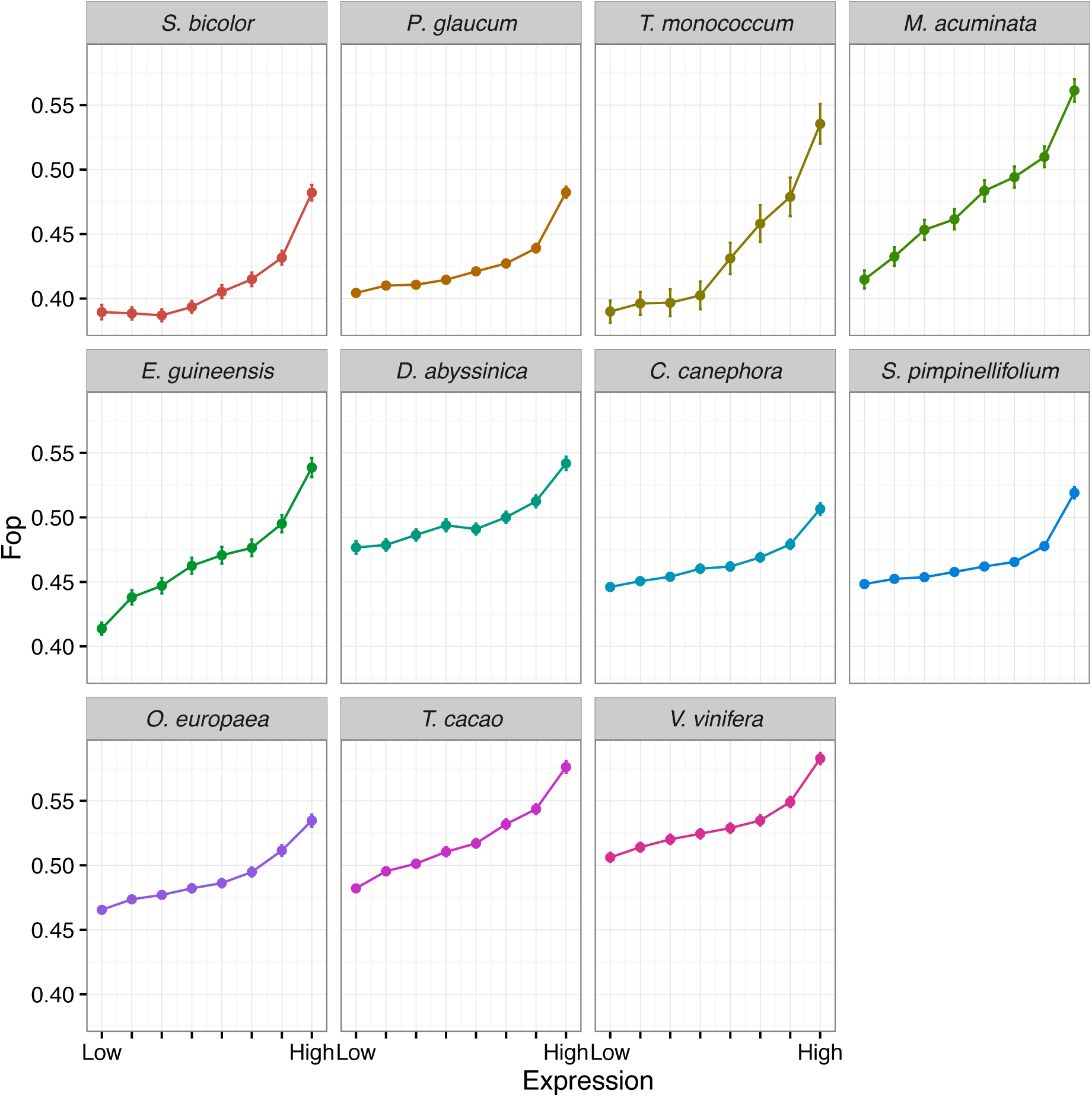
Relationship between the frequency of optimal codons (FOP) and expression in the 11 studied species. For each species, genes have been split into eight categories of expression (based on RPKM) of same size and the mean FOP for each category is plotted with its 95% confidence interval.

### Selective-like evolutionary forces affect base composition

To determine which forces affect variation in base composition and codon usage among species, we first evaluated whether base composition at synonymous sites was at mutation-drift equilibrium. Glémin et al. [33] showed that the asymmetry of the distribution of non-polarized GC allele frequencies (measured by the skewness coefficient) is a robust test of this equilibrium. This statistic is not affected by possible polarization errors (see later for more on polarization errors). Null skewness is expected under equilibrium whereas negative (resp. positive) skewness means higher (resp. lower) GC content than expected under mutation-drift equilibrium. The same rationale can be applied to codon frequencies. We found that GC content and the frequency of preferred codons are significantly higher than predicted by mutational effects in all species, with the exception of coffee, which interestingly showed a lower GC content than expected under mutation-drift balance (Table 3).

**Table 3:**
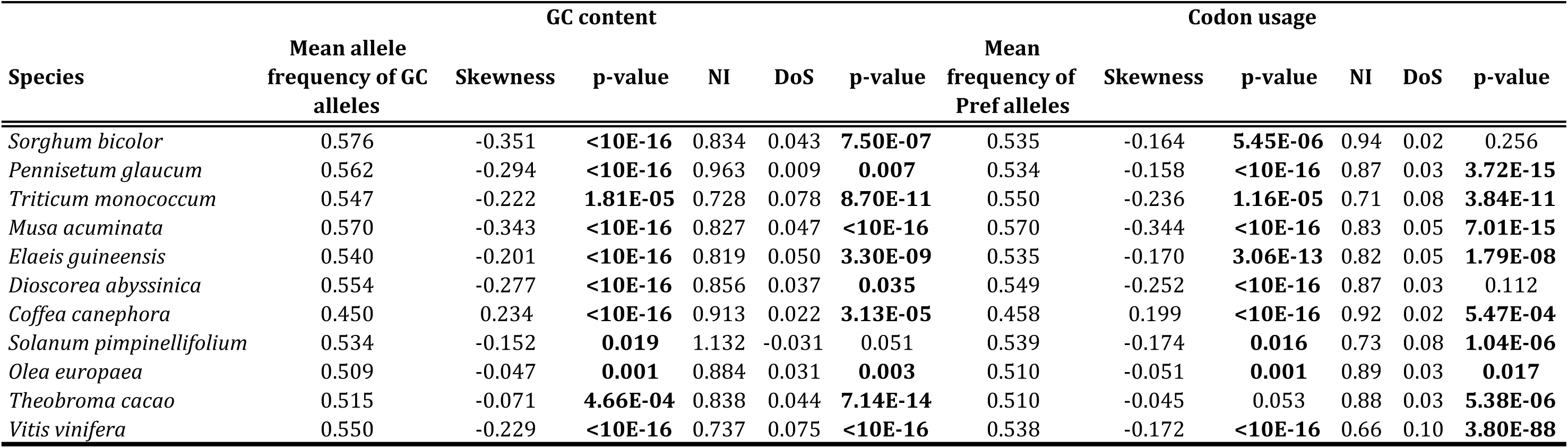
Skewness, NI and DoS statistics for GC content and codon usage.

As base composition equilibrates slowly under mutation pressure [28], non-equilibrium conditions could be due to long-term changes in mutational patterns. To test further whether selective-like forces can explain the excess of GC and preferred codons, we developed a modified MacDonald Kreitman test [46] comparing W→S (resp. U→P) to S→W (resp. P→U) polymorphic and divergent sites (Material & Methods and Text S1). SNPs and fixed mutations (substitutions) were polarized by parsimony using two outgroup taxa for each focal species. We built contingency tables by counting the number of polymorphic or divergent sites for each of the two mutational categories. From these contingency tables, we computed neutrality, NI, [47] and direction of selection, DoS, [48] indices. In the case of selective-like forces favouring the fixation of W→S or U→P mutation, we expect NI values to be lower than 1 and DoS values to be positive. P-values were computed from a Chi-squared test on the contingency tables. Results show that NI is lower than 1 and DoS positive in all species except *S. pimpinellifolium* (Table 3), indicating that selective-like forces drive the fixation of GC and preferred codon alleles. In *P. glaucum*, although significant, the departure from the neutral expectation for GC content is minute, which can be explained by very weak gBGC but also by a recent increase in its intensity (see results below and Text S1). Overall, this analysis showed that in most species selective-like forces tend to drive base and codon composition away from their mutational equilibrium. Selection and gBGC are the two known alternatives which effects have to be distinguished.

### Disentangling gBGC and SCU?

Although they may have different mechanistic causes and biological consequences, selection and gBGC leave similar evolutionary footprints and are not easy to disentangle, especially in species where most preferred codons end in G or C (Table 2). We first applied correlative approaches to try to disentangle both processes. Then we tried to quantify their respective intensities.

Under the SCU hypothesis, departure from neutrality should be stronger for highly expressed genes and/or genes with strongly biased codon composition. Under the gBGC hypothesis, departure from neutrality should increase with recombination rates. However, recombination data is not available in our datasets. As gBGC leads to an increase in GC content, departure from neutrality should thus also increases with GC content. We split synonymous SNPs and substitutions into eight groups of same size according to their GC3 or their gene expression level (measured by the mean RPKM values across all individuals of a given population), and computed the NI and DoS indices for each category based on W/S or U/P changes. For all species except *D. abyssinica* and *S. bicolor*, we found a strong positive (resp. negative) correlation between GC3 and DoS (resp. NI), indicating a stronger bias in favour of S alleles in GC-rich genes (Figure 4). In contrast, the relationship between expression level and DoS or NI measured on codon usage is weaker, with more variable and on average lower correlation coefficients (Figure 4). These results tend to point out gBGC as a stronger force than SCU affecting synonymous variations in our datasets.

**Figure 4:**
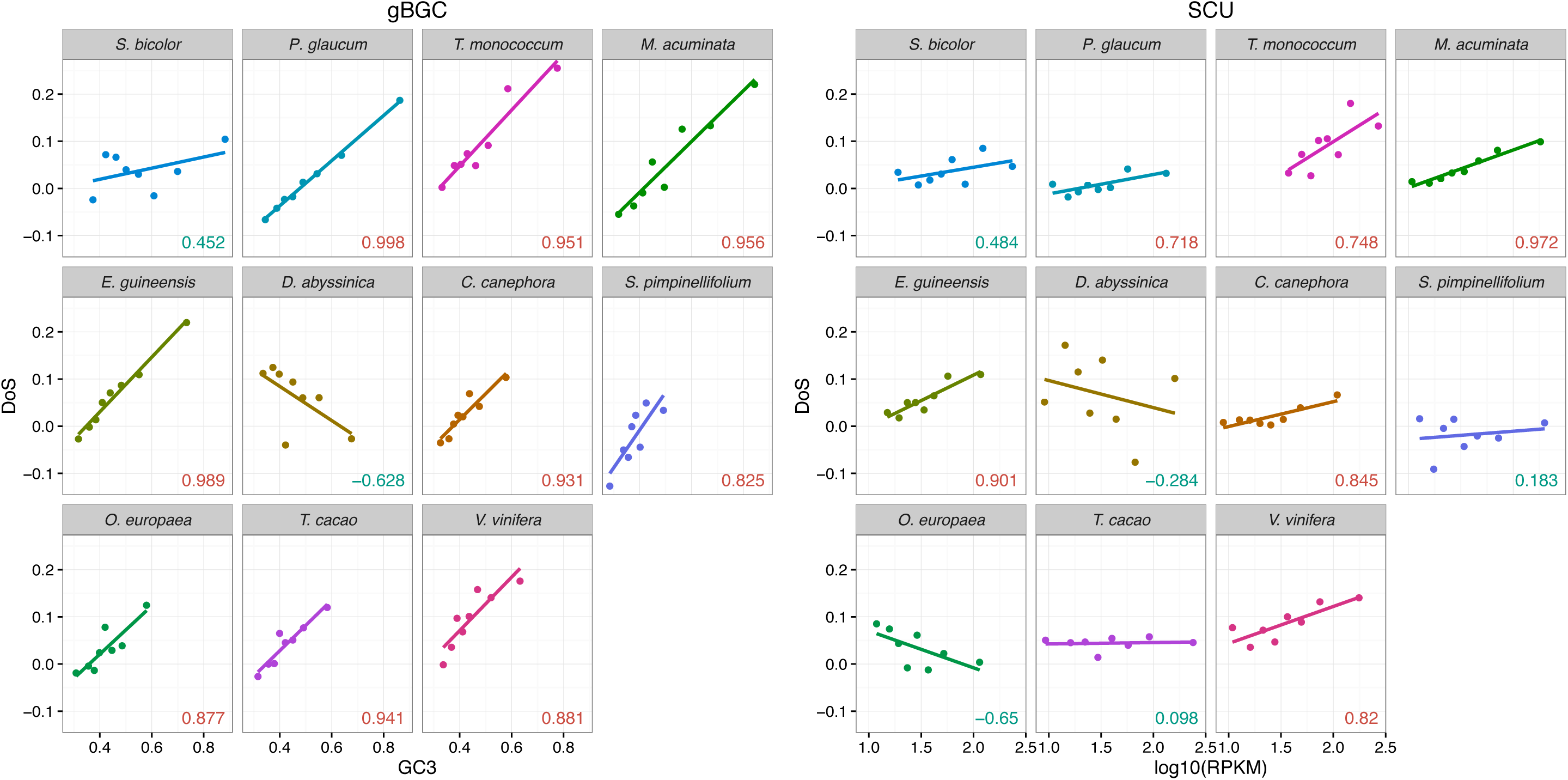
DoS statistics as a function of GC3 and expression level. Correlation between GC3 and DoS computed on WS changes (left panel) or between expression level (measured through RPKM) and DoS computed on UP changes (wright). Pearson correlation coefficients are given for each species (red: significant at the 5% level, blue nonK significant).

We then split our datasets into four independent categories based on two GC3 groups crossed by two expression level groups to test which factor has the strongest effect on the bias towards S or P alleles. The rationale is that SCU should make the bias towards P alleles increase with gene expression independently of GC3. On the other hand, gBGC should increase the bias towards S alleles with GC3 independently of gene expression. We found that DoS clearly increases with GC3 in all species for both lowly and highly expressed genes, except in *D. abyssinica* and *S. bicolor* where it decreases for lowly expressed genes. On the other hand, the effect of expression on DoS is inconsistent or only weak in most species (Figure 5). These results confirm that the effect of gBGC appears stronger than the effect of SCU.

**Figure 5:**
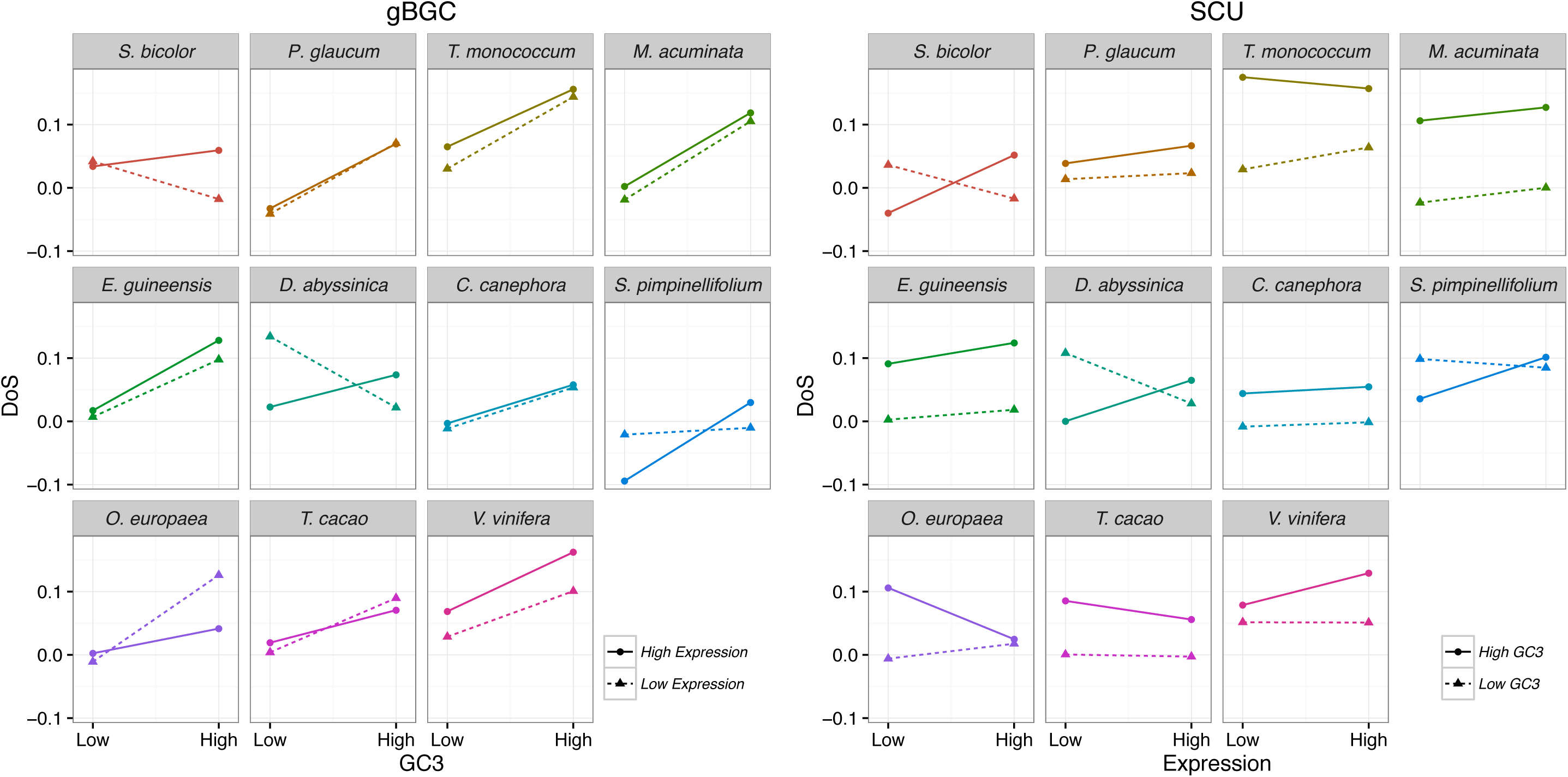
Combined effect of GC3 and expression level on DoS statistics. The DoS statistics was computed on W/S (gBGC) or U/P (SCU) changes for four gene categories: GCKrich and highly expressed, GCKrich and lowly expressed, GCKpoor and highly expressed, GCKpoor and lowly expressed.

### Estimation of gBGC/SCU intensity and mutational bias

To evaluate further the forces affecting base composition we estimated the intensity of selection (*S* = 4*N*_*e*_*s*) and gBGC (*B* = 4*N*_*e*_*b*) from site frequency spectra (SFS). SFS for all species are given in Figure S3. We used the method recently developed by Glémin et al. [33] that takes SNP polarization errors into account, which avoids observing spurious signature of selection or gBGC. As mentioned above, the observed pattern in *P. glaucum* (excess of GC content but almost no departure from neutrality according to the NI and DoS indices, see Table 3) suggests a recent change in the intensity of selection and/or gBGC. Also, transition to selfing, which usually can be very recent in plants [49], could have effectively shut down gBGC in the recent past due to a deficit in heterozygous positions. To capture these possible changes of fixation bias through time, we extended the model of [33] by combining frequency spectra and divergence estimates as summarized on Figure 6 (and see Text S2 for full details). Divergence is determined by both mutation and selection/gBGC so it is not possible to disentangle these two factors from the divergence data alone. However, if we assume constant and identical mutation bias at the polymorphism and the divergence level, there are enough degrees of freedom to fit an additional *S* or *B* parameter. Thus, we assumed a single mutation bias but two different selection/gBGC intensities, one fitted on polymorphism data and the other on divergence. We evaluated the statistical significance of the shift in intensity by a likelihood ratio test with the model where the two intensities are equal (*i.e.* no change over time). Simulations showed that not taking polarization errors into account can bias selection/gBGC estimates as already shown in [33] and also leads to spurious detection of changes in selection/gBGC intensities (Text S2). Simulations also showed that the estimated differences between the two intensities are often underestimated. This is expected as *B* values estimated in the model correspond to averages over the conditions that mutations have experienced during their lifetime (drift and gBGC/selection intensities), so it depends on when changes occurred. Overall, the test of heterogeneity of selection/gBGC is a conservative approach.

**Figure 6:**
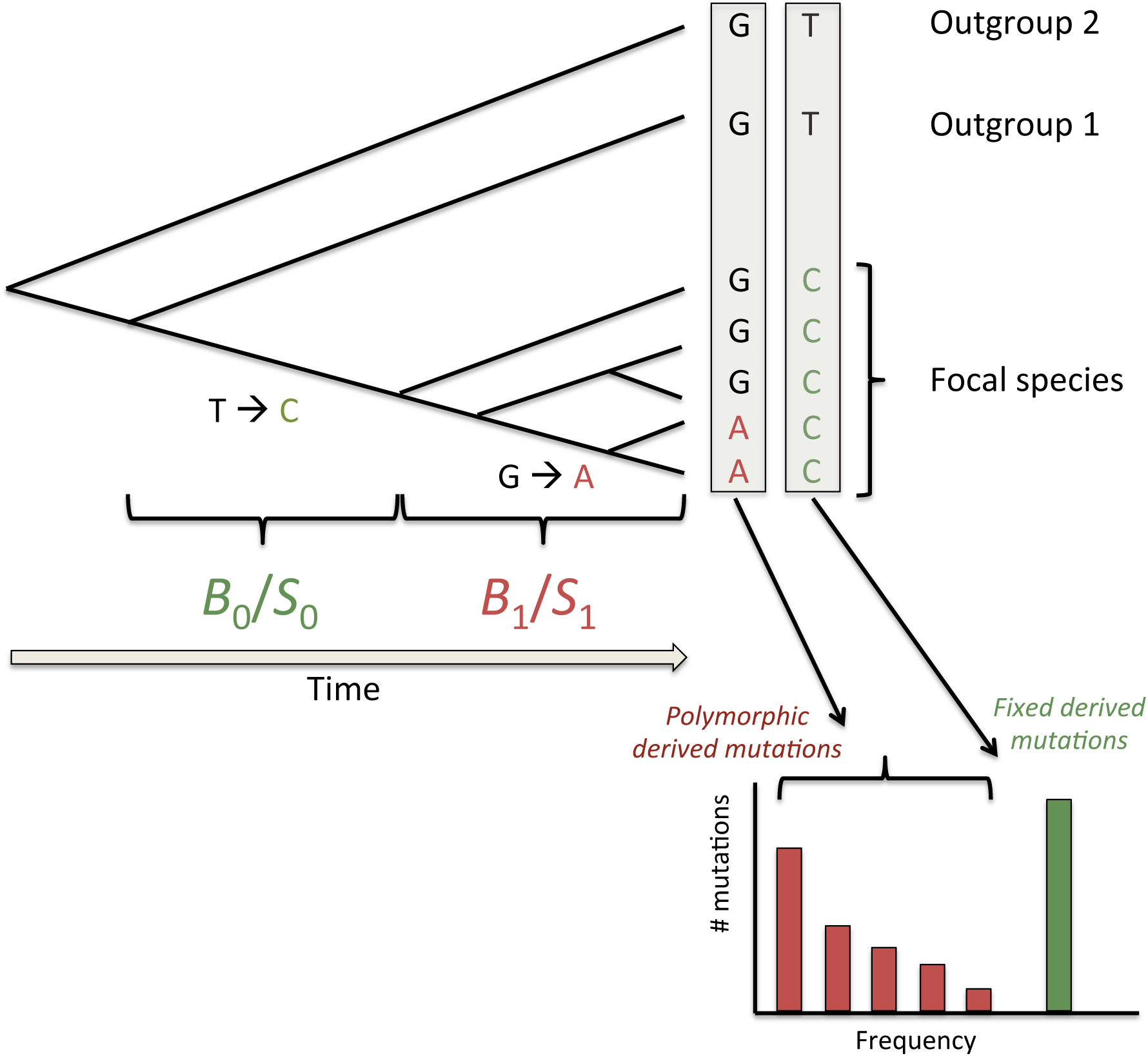
schematic presentation of the method to estimate recent and ancestral gBGC or SCU. In addition to polymorphic derived mutations used to infer recent gBGC or selection (*B*_1_/*S*_1_) as in [33] we also consider substitutions (*i.e.* fixed derived mutations) on the branch leading to the focal species. Each box corresponds to a site position in a sequence alignment. Both kinds of mutations are polarized with the two same outgroups and are thus sensitive to the same probability of polarization error. We assume that gBGC and selection may have change so that fixed mutations may have undergo a different intensity. Note that these two *B* or *S* values correspond to average of potentially more complex variations over the two periods.

We applied the method to the total frequency spectra, either for W/S or U/P polymorphisms and substitutions. In all species, significant (at the 5% level) gBGC or SCU were detected but at low intensity (*B* or *S* < 1, Table 4). In four species (*P. glaucum, E. guineensis, D. abyssinica* and *V. vinifera*) we found significant differences between ancestral and recent intensities for gBGC and/or SCU. In particular, the recent increase in gBGC/SCU in *P. glaucum* explains why NI is very close to one (or DoS close to zero) (see above and Text S1). On average, Monocots, especially Commelinids species tend to exhibit stronger gBGC than Eudicots and *B* tends to increase with mean GC3, but no relationship is significant with only 11 species when either *B*0 or *B*1 are used. However, using the constant *B* estimates (Table S4), weakly significant relationships were found for the difference between Commelinids and other species (Wilcoxon test: p-value = 0.0519) and the correlation between *B* and GC3 (ρ_Spearman_ = 0.691, p-value = 0.023). No significant relationship was found for SCU.

**Table 4:** Separated estimations of recent and ancestral gBGC (*B* = 4*N*_*e*_*b*) and SCU (*S* = 4*N*_*e*_*s*)

| Species | lambda | $4N_e b$ ancestral | $4N_e b$ recent | gBGC | | |
| --- | --- | --- | --- | --- | --- | --- |
|  |  |  |  | p-value ancestral = 0 | p-value recent = 0 | p-value recent = ancestral |
| *Sorghum bicolor* | 1.61 [1.51 - 2.69] | 0.378 [0.290 - 0.516] | 0.078 [-0.492 - 0.739] | **2.73E-14** | 0.758 | 0.189 |
| *Pennisetum glaucum* | 1.73 [1.69 - 1.83] | 0.224 [0.189 - 0.261] | 0.524 [0.383 - 0.661] | **<10E-16** | **1.15E-13** | **2.18E-06** |
| *Triticum monococcum* | 1.99 [1.67 - 2.25] | 0.448 [0.269 - 0.613] | -0.008 [-0.824 - 0.691] | **1.39E-05** | 0.985 | 0.164 |
| *Musa acuminata* | 1.71 [1.66 - 1.80] | 0.313 [0.253 - 0.370] | 0.397 [0.234 - 0.546] | **<10E-16** | **2.68E-06** | 0.343 |
| *Elaeis guineensis* | 1.84 [1.77 - 1.93] | 0.328 [0.267 - 0.400] | 0.516 [0.328 - 0.702] | **<10E-16** | **1.76E-07** | **0.034** |
| *Dioscorea abyssinica* | 2.20 [2.10 - 2.47] | 1.171 [0.127 - 4.067] | 0.008 [-0.221 - 0.264] | **0.032** | 0.949 | 0.072 |
| *Coffea canephora* | 1.05 [1.02 - 1.10] | 0.154 [0.110 - 0.202] | 0.243 [0.113 - 0.366] | **9.47E-11** | **3.77E-04** | 0.171 |
| *Solanum pimpinellifolium* | 2.05 [1.74 - 2.63] | 0.114 [-0.057 - 0.392] | 0.759 [-0.491 - 3.785] | 0.215 | 0.153 | 0.193 |
| *Olea europaea* | 1.58 [1.53 - 1.64] | 0.167 [0.080 - 0.268] | 0.031 [-0.127 - 0.168] | **<10E-16** | 0.687 | 0.132 |
| *Theobroma cacao* | 1.67 [1.59 - 1.74] | 0.316 [0.258 - 0.377] | 0.465 [0.222 - 0.683] | **<10E-16** | **6.54E-05** | 0.135 |
| *Vitis vinifera* | 2.15 [2.08 - 2.22] | 0.360 [0.318 - 0.413] | 0.024 [-0.101 - 0.153] | **<10E-16** | 0.71 | **1.55E-08** |

| Species | lambda | $4N_e s$ ancestral | $4N_e s$ recent | SCU | | |
| --- | --- | --- | --- | --- | --- | --- |
|  |  |  |  | p-value ancestral = 0 | p-value recent = 0 | p-value recent = ancestral |
| *Sorghum bicolor* | 2.04 [1.70 - 2.47] | 0.139 [0.023 - 0.260] | 0.439 [-0.251 - 1.083] | **0.010** | 0.143 | 0.341 |
| *Pennisetum glaucum* | 1.76 [1.70 - 1.87] | 0.181 [0.137 - 0.226] | 0.126 [-0.062 - 0.289] | **2.33E-15** | 0.165 | 0.484 |
| *Triticum monococcum* | 2.84 [2.33 - 3.31] | 0.534 [0.353 - 0.718] | 0.236 [-0.610 - 1.029] | **1.14E-06** | 0.581 | 0.409 |
| *Musa acuminata* | 2.02 [1.96 - 2.15] | 0.315 [0.256 - 0.362] | 0.392 [0.221 - 0.553] | **<10E-16** | **5.21E-06** | 0.394 |
| *Elaeis guineensis* | 1.58 [1.50 - 1.66] | 0.324 [0.233 - 0.396] | 0.512 [0.322 - 0.704] | **3.00E-15** | **6.51E-07** | **0.043** |
| *Dioscorea abyssinica* | 1.68 [1.39 - 1.74] | 1.909 [0.306 - 9.994] | -0.101 [-0.311 - 0.135] | **0.023** | 0.470 | **0.037** |
| *Coffea canephora* | 0.89 [0.86 - 0.95] | 0.148 [0.079 - 0.197] | 0.196 [0.039 - 0.330] | **5.91E-08** | **0.012** | 0.515 |
| *Solanum pimpinellifolium* | 1.56 [1.32 - 2.05] | 0.465 [0.270 - 0.857] | 0.566 [-0.567 - 3.900] | **3.39E-06** | 0.285 | 0.834 |
| *Olea europaea* | 1.18 [1.13 - 1.22] | 0.148 [0.040 - 0.241] | 0.025 [-0.162 - 0.186] | **0.004** | 0.772 | 0.214 |
| *Theobroma cacao* | 1.09 [1.02 - 1.16] | 0.245 [0.167 - 0.339] | 0.397 [0.107 - 0.673] | **2.85E-11** | **3.00E-03** | 0.185 |
| *Vitis vinifera* | 1.26 [1.22 - 1.32] | 0.470 [0.421 - 0.525] | 0.118 [-0.028 - 0.258] | **<10E-16** | 0.103 | **7.09E-08** |

As the two processes are entangled, it is difficult to properly and separately estimate their respective intensities. To do so, we developed a second extension of the method of [33]. Combining the two processes, nine kinds of mutations can occur (see Text S2). By assuming that selection and gBGC act additively, it is in theory possible to estimate separately the two effects. We fit a general model to the nine SFS and the nine substitution counts, with a constant mutation bias, two *B* and two *S* values. The details of the model are reported in Text S2. Simulations show that the method can efficiently estimate both gBGC and SCU but tends to slightly underestimate recent gBGC and overestimate recent SCU (Text S2). When the distributions of SNPs and substitutions are highly unbalanced (typically S/P and W/U states are confounded), it is more difficult to detect both effects with a significant level (Text S2). For both selection and gBGC and both ancestral and recent periods, we either fixed the value to 0 or let it be freely estimated, leading to 16 different models. For each species, the best model according to AIC criteria (see methods) is given in Table 5 while all results are given in Table S5. In six species the model with only gBGC is the best one, this could also include *M. acuminata* where it was not possible to disentangle between gBGC and SCU. For three species, the best model includes both gBGC and SCU and only *S. pimpinellifolium* appears to be affected by SCU but not gBGC. Overall, this confirms that synonymous sites are mostly affected by gBGC in plant species and that SCU only plays a minor role. This result is expected to be robust and conservative because simulations suggest that SCU is slightly more easily detected than gBGC.

**Table 5:** Best model for the joined estimations of recent and ancestral gBGC (*B* = 4*N*_*e*_*b*) and SCU (*S* = 4*N*_*e*_*s*). For *Musa acuminata* the two best models with very close AIC values are given.

| Species | $4N_e b$ ancestral | $4N_e b$ recent | $4N_e s$ ancestral | $4N_e s$ recent |
| --- | --- | --- | --- | --- |
| *Sorghum bicolor* | 0.439 [0.334 - 0.525] | 0 | 0 | 0 |
| *Pennisetum glaucum* | 0.218 [0.182 - 0.253] | 0.561 [0.393 - 0.689] | 0.139 [0.106 - 0.175] | 0 |
| *Triticum monococcum* | 0.264 [0.042 - 0.443] | 0 | 0.247 [0.027 - 0.468] | 0 |
| *Musa acuminata 1* | 0.312 [0.281 - 0.395] | 0.394 [0.241 - 0.580] | 0 | 0 |
| *Musa acuminata 2* | 0 | 0 | 0.317 [0.284 - 0.400] | 0.398 [0.176 - 0.540] |
| *Elaeis guineensis* | 0.329 [0.241 - 0.383] | 0.517 [0.234 - 0.744] | 0 | 0 |
| *Dioscorea abyssinica* | 1.256 [0.564 - 2.202] | 0 | 0 | 0 |
| *Coffea canephora* | 0.154 [0.119 - 0.227] | 0.244 [0.070 - 0.361] | 0 | 0 |
| *Solanum pimpinellifolium* | 0 | 0 | 0.459 [0.311 - 0.603] | 0 |
| *Olea europaea* | 0.168 [0.074 - 0.250] | 0 | 0 | 0 |
| *Theobroma cacao* | 0.318 [0.241 - 0.383] | 0.474 [0.234 - 0.744] | 0 | 0 |
| *Vitis vinifera* | 0.256 [0.216 - 0.295] | 0 | 0.380 [0.323 - 0.439] | 0 |

This method also allows us to estimate mutation bias. As already observed in most species, mutation is biased towards AT alleles, with a bias slightly ranging from 1.6 to 2.2 (Table 4), which is of the same order as what was found in humans [33,50]. Interestingly, *C. canephora* is again an exception with almost no mutational bias (λ = 1.05).

### Variation along genes

So far, we considered either global effects at the transcriptome scale or variations among genes belonging to different categories. However, most plant species exhibit a more or less pronounced gradient in base composition from 5’ to 3’ [1], which is strongly linked to exon-intron structure [32]. In particular, in some species the first exon is much GC-richer than other exons. Moreover, it has been proposed that this gradient could be due to a gBGC gradient associated with a recombination gradient [28]. To quantitatively test this hypothesis, we separated SNPs and fixed derived mutations as a function of their position along genes. The best choice would have been to split them according to exon ranking [32]. However, as exon annotation is lacking (or imprecise) for most species in our datasets, we split contigs into two sets: the first 252 base pairs, corresponding to the median length of the first exon in *Arabidopsis*, banana and rice (Gramene database [51]), used as a proxy for the first exon, and the rest of the contig. We then estimated *B* on these two sets of contigs.

For all species except *D. abyssinica* and *S. pimpinellifolium*, the ancestral *B* is higher in the first part than in the rest of contigs but the signature is less clear for recent *B* as far less values are significant though ancestral and recent *B* are not significantly different in most species (Table S6). To illustrate the global pattern, Figure 7 shows average gBGC gradients for all species, *i.e.* assuming the same ancestral and recent *B* values. Interestingly, while there is no clear taxonomic effect on global gBGC estimates (Table 4), there is a sharp difference between Commelinid species and the others for the first part of contigs (Wilcoxon test p-value = 0.030, Figure 7C), in agreement with the strong 5’ – 3’ GC gradient observed in these species [1,2]. Moreover, *B* values and GC3 tends to be more positively correlated on the first part of contigs (ρ_Spearman_ = 0.591, p-value = 0.061) than in the rest (ρ_Spearman_ = 0.382, p-value = 0.248). These analyses were performed on all contigs but some of them do not start by a start codon. We restricted the analyses to the subset of contigs starting by a start codon and we found very similar results with stronger statistical supports (Table S6 and Figure S4). In line with previous results showing that first exons contribute to most of the variation in GC content among species [2,28,32], these results show that species also mostly differ in their gBGC intensities in the first part of genes.

**Figure 7:**
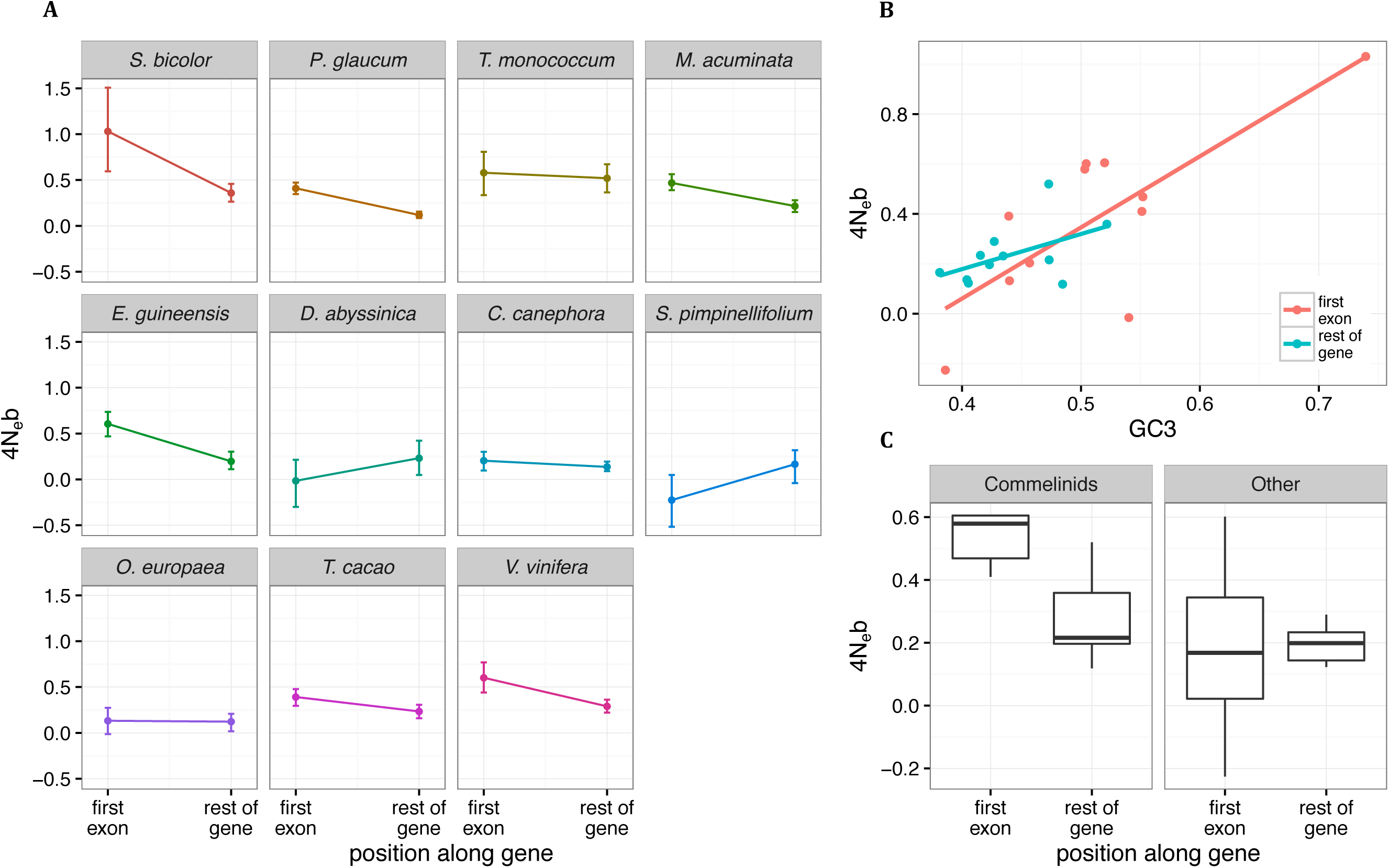
GC3 and gBGC gradients along genes.

## Discussion

### Selective-like evolution of synonymous variations in plant genomes

It has already been shown that base composition in grass genomes is not at mutation-drift equilibrium with both gBGC and selection increasing GC content despite mutational bias toward A/T [26]. Our results demonstrate that even in GC-poor genomes base composition is not at mutation-drift equilibrium, implying that selective-like forces are widespread in all the 11 plant species we studied. In all species, either the skewness and/or the DoS/NI statistics show evidence of departure from equilibrium and purely neutral evolution (Table 3). All species except *C. canephora* have higher GC content than predicted by mutational effect alone, which could be explained by a mutation/gBGC (or selection)/drift balance.

The case of *C. canephora* remains intriguing. Mutation seems not to be biased towards AT as observed in all mutation accumulation experiments [reviewed in 52] and through indirect methods [53]. So far, GC biased mutation has only been observed in the bacteria *Burkholderia cenocepacia* [52]. However, despite no apparent or very weak AT mutational bias and evidence of both recent and ancestral gBGC (Table 4), GC content is rather low (GC3 = 0.42, Table 2) and lower than expected under mutation pressure alone (1/(1+λ) = 0.49) as revealed by the positive skewness statistics (Table 3). Preferred codons mostly end in G or C (Table 2) so that SCU is not a possible explanation for this low GC content. Rather, a recent change in mutation bias is a more probable explanation. Using *B*_0_ = 0.154 or *B*_1_ = 0.243 (Table 4), a mutational bias of 1.61 or 1.76 would be necessary to reach the observed GC3 (= 0.42). Such values are in the same range as observed for the other species. So further investigation of mutational patterns in this species would be useful to understand better its intriguing base composition pattern.

Beyond departure from equilibrium, comparison of ancestral and recent gBGC or selection also reveals the dynamic nature of forces affecting base composition. At least four species (*P. glaucum, E. guineensis, D. abyssinica* and *V. vinifera*) show evidence of significant change in gBGC and/or SCU intensity over time (Table 4). If we consider the first part of genes only, changes also occurred in *M. acuminata* and *T. cacao* (Table S6). Moreover, our method is conservative (see Text S2) so we may have missed variations in other species. Changes occurred in both directions. In the three selfing or mixed mating species (*S. pimpinellifolium, T. monococcum*, and *S. bicolor*) the ancestral gBGC or SCU intensity is significantly positive but the recent one is null. This is supported by the rather recent evolution of selfing in these species, which nullifies the effect of gBGC through the increase in homozygosity levels and reduces the efficacy of selection [54]. In other species, gBGC or SCU have increased (e.g. *P. glaucum*) or decreased (e.g. *V. vinifera*). Recalling that *B* = 4*N*_*e*_*rb*_0_ (see introduction), this could be explained by changes in effective population size (*N*_*e*_) recombination rate (*r*), gBGC intensity per recombination event (*b*_0_) and also conversion tract length, which might also vary among species [55]. To date, we know nothing about the stability of *b*_0_ and how fast it can evolve. In some species, such as mammals, recombination can evolve very rapidly, at least at the hotspot scale [56] but it can be more stable in other species like in birds [57], yeast [58] or maize [59]. Moreover, we average gBGC over the whole transcriptome so recent genome-scale changes in recombination should be necessary to explain changes in *B*. Although recent changes in *r* and *b*_0_ are possible, changes in effective population size over time appears to be the most likely explanation.

Selective-like evolution and non-equilibrium conditions can have practical impacts on several genomic analyses. First, gBGC can lead to spurious signatures of positive selection [60], significantly increasing the rate of false positive in genome scan approaches in mammals [61]. This problem should also be taken into account in plant genomes, even in GC-poor ones. Second, SCU/gBGC and non-stationary evolution, due for instance to changes in population size, can strongly affect the estimation of the rate of adaptive evolution through McDonald-Kreitman approaches, especially at high GC content [62]. In species far from equilibrium such as Commelinids, it should be an issue to consider.

## gBGC, SCU or both?

### Technical issues

We found clear evidences that base composition evolution is not driven only by mutation. However, it was more difficult to distinguish gBGC from SCU because we only used coding regions in our study. Unfortunately, we were not able to use 5’ or 3’ flanking regions to compare them with synonymous coding positions. These flanking regions were too short and of lower sequencing coverage and quality: they were not frequently sequenced and corresponded to sequence ends. Comparison with introns or non-coding regions would be helpful in the future to confirm our findings, as it was done in rice [26] or maize [27]. To bypass this problem, we developed a new method that jointly estimates gBGC and SCU and allows testing which processes are significant. However, the two processes are especially difficult to distinguish in species where most preferred codons end in G or C, such as *M. acuminata* and *T. monococcum* (Table. 2 and 5 and Text S2). However, simulations suggest that weaker SCU than gBGC could be estimated even for a highly unbalanced dataset (at least ancestral SCU, see Text S2). Finally, correlative approaches with GC content and expression can also help distinguishing the two processes. Overall, although each individual result (species-specific and or approach-specific) can be insufficiently conclusive, they collectively point towards the general conclusion of a major contribution of gBCG over SCU to explain synonymous variation in the studied plant species.

### gBGC seems to predominate

The combination of our different results clearly shows that gBGC prevails over SCU in the studied plants. While signatures of gBGC were detected in all species but *S. pimpinellifolium*, SCU was detected only in four or five species (Table 5). However, in these species, the change in NI/DoS with expression is consistent with SCU only in *P. glaucum* (Figure 4). These poorly supported results do not necessarily mean that SCU is not active. Indeed, we were able to defined preferred codons in all our species, and Fop increases with expression level in all of them (Figure 2). However, changes in Fop with expression are moderate to low (15% to 5%) and on average lower to what was observed in *Drosophila* (15%) or *Caenorhabditis* (25%), but slightly higher than *Arabidopsis* (5%) [44]. Thus, SCU is likely active but at a level too low to be detected by our methodology in some species, especially because gBGC masks its effect. A larger dataset (increasing both the number of SNPs and of individuals) would probably be necessary to properly estimate SCU in the presence of gBGC, especially when the most preferred codons end with G or C. It should be noted that in *P. glaucum*, one of the species where SCU was quite confidently detected, a high number of SNPs and a rather equilibrated patterns of codon preference were identified.

### Coevolution between GC and codon usage?

The difficulty in distinguishing gBGC and SCU also raises the question of the interaction between these two processes. The predominance of GC ending preferred codons has also been observed in many bacteria [63]. The bias towards GC ending preferred codons increases with genomic GC content, with species having a GC content higher than 40% being strongly biased towards GC preference [63]. The classical Bulmer’s model of coevolution between preferred codons and tRNA predicts a match between the frequency of tRNAs and preferred codons with two equivalent stable states (either AT or GC preference), and so does not explain the observed bias in preference [64]. However, our results are compatible with a modified version of this model in which an external force on base composition is introduced [65]. gBGC could act as such a force. By increasing GC content, gBGC could disrupt the co-evolutionary equilibrium between preferred codons and tRNAs abundance towards a higher level of GC preference. This would in turn leads to the confounding effects of gBGC and SCU.

### GC content gradient and the gBGC hypothesis

We detected gBGC in all but one species but its intensity is rather weak (Table. 4 and 5 and S4 and S5), of the same order to what was estimated in humans [33] but lower than in other mammals [34], maize [66], and particularly honey bee [37]. Low values can be explained by the fact that we only estimated average *B* values. In many plants studied so far, recombination was found to be heterogeneous along chromosomes [e.g. 31] and locally occurring in hotspots [e.g. 29,30,59], so that gBGC can be locally much higher than average estimates. However we did not apply the hotspot model proposed by [33] because it behaves poorly when not constrained by additional information on hotspot structure, which we lack in the species studied here. In addition, recombination hotspots are preferentially located outside genes, especially in 5’ upstream regions (and 3’ downstream regions to a lesser extent) [29,30,31]. As we estimated gBGC intensities within coding regions, this can also explain why we only estimated rather weak *B* values.

A consequence of this specific recombination hotspot location is the induction of a 5’ – 3’ recombination gradient along genes (or more generally an exterior to interior gradient if also considering downstream location) [29,30]. Recently, it has been proposed that this recombination gradient could explain the 5’ – 3’ gradient observed in grasses and more generally in many plant species [28]. We tested this model by looking at signatures of gBGC along contigs in our datasets. In agreement with this model, we found stronger gBGC signatures at the 5’ end of contigs compared to the rest of contigs in most of our species (Figure 7). The fact that we observed this gBGC gradient in both Eudicots and Monocots suggests that all these species share the same meiotic recombination structure with preferential location of recombination in upstream regions of gene, which was hypothesized to be the ancestral mode of recombination location in Eukaryotes [67].

Glémin et al. [28] also proposed that changes in the steepness of the recombination/gBGC gradient could explain variation in GC content distributions among species, from unimodal GC-poor to bimodal GC-rich distributions. Alternatively, if gradients are stable among species, changes in gene structure, especially the number of short mono-exonic genes and the distribution of length of first introns, could also generate variations in GC content distribution [28,32]. Here we found that, in the first part of genes, gBGC is the highest in Commelinid species, which exhibit the richest and most heterogeneous GC content distributions (Figure 7). This result parallels the sharp difference in GC content in first exons between rice and *Arabidopsis* whereas the centres of genes have a very similar base composition [32]. Our results support the hypothesis that genic base composition in GC-rich and heterogeneous genomes has been driven by high gBGC/recombination gradients. As GC-content bimodality is likely ancestral to monocot species and has been lost several times later [2], our results suggest that an increase in gBGC and or recombination rates occurred at the origin of the Monocot lineage.

## Conclusion

Overall, we show that selection on codon usage only plays a minor role in shaping base composition evolution at synonymous sites in plant genomes and that gBGC is the main driving force. Our study comes along an increasing number of results showing that gBGC is at work in many organisms. Plants are no exception. If, as we suggest, gBGC is the main contributor to base composition variation among plant species, it shifts the question towards understanding why gBGC may vary between species and more generally why gBGC evolved. Our results also imply that gBGC should be taken into account when analysing plant genomes, especially GC-rich ones. Typically, claims of adaptive significance of variation in GC content should be viewed with caution and properly tested against the “extended null hypothesis” of molecular evolution including the possible effect of gBGC [60].

## Materials & Methods

### Dataset

We focused our study of synonymous variations on 11 species spread across the Angiosperm phylogeny with contrasted base composition and mating systems, *Coffea canephora, Olea europaea, Solanum pimpinellifolium, Theobroma cacao, Vitis vinifera, Dioscorea abyssinica, Elaeis guineensis, Musa acuminata, Pennisetum glaucum, Sorghum bicolor* and *Triticum monococcum*. A phylogeny of these species is shown in Figure 1. For practical reasons, we chose diploid cultivated species but focused our analysis on wild populations except in *Elaeis guineensis* where domestication is very recent and limited (19th century [41]). Using the same methodology as [42], we sequenced for each species the transcriptome of ten individuals (12 in the case of *C. canephora* and *V. vinifera*, nine in the case of *S. bicolor* and five in the case of *D. abyssinica*) plus two individuals coming from two outgroup species, using RNA-seq (See Text S3 for details). After cleaning, reads were either mapped on the transcriptome extracted from the reference genome (when available, see Table 1) or on the de novo transcriptome of each species (including outgroups) obtained from [42]. For *C. canephora* and its outgroups, no transcriptome was available. We thus applied the same methodology and pipeline as in [42] to assemble and annotate contigs. For banana, *M. acuminata*, Robusta coffee tree, *C. canephora*, and for the outgroup *Phoenix dactylifera*, genome sequences were available but the quality of mapping was rather poor because of problem of definition of exon/intron boundaries. We thus prefer to assemble a new transcriptome from our data using the same protocol. Details of the assemblies for all species are given in Table S2. Details of data processing are provided in Text S4. Only contigs with at least one mapped read for each individual was kept for further analysis. Expression levels for each individual in each contig were computed as RPKM values (*i.e.* the number of Reads per Kilobase per Millions mapped reads). We called genotypes and filtered out paralogs for each species individually using the *read2snp* software [43] (see Text S4 for details). Genotypes were called when the coverage was at least 10x and the posterior probability of the genotype higher than 0.95. Otherwise, the genotype of the individual was considered as missing data. Orthology between focal and outgroup individuals was determined by best reciprocal blast hit. Finally, we aligned orthologous contigs (focal and outgroup individuals) sequences using MACSE [68].

### SNPs detection and polarization

We scanned contig alignments in each focal species for polymorphic sites. We only considered gapless sites for which all focal individuals were genotyped. Only bi-allelic SNPs were considered. In the highly selfing *T. monococcum*, the deficit in heterozygous sites can lead to abnormal site frequency spectra that are difficult to analyse. We thus used an allele sampling procedure that effectively divides the number of chromosomes by two by merging together homologous chromosomes in each individual. For heterozygous sites, one allele was randomly chosen. For the mixed mating *S. bicolor* and *S. pimpinellifolium*, we used the full SFSs.

SNPs were polarized using parsimony by comparing alleles in focal individuals to orthologous positions in outgroups. For each polymorphic site, the ancestral allele was inferred to be the one identical to both outgroup species, while the other allele was inferred to be derived. All polarized SNPs are marked ancestral → derived for the remainder of the paper. A and T bases were grouped together as W (for weak) while G and C bases were grouped together as S (for strong). We thus classified mutations as W→S, S→W or neutral with respect to gBGC (S←→S or W←→W).

### SNPs and preferred codons

In each species, preferred (P) and un-preferred (U) codons were defined using the ΔRSCU method [44]. In each contig, we computed for each codon its RSCU value, or relative frequency (i.e. its number of occurrences in a contig normalized by the number of occurrences of its amino-acid in the same contig). Contigs were divided into eight groups of identical size based on their expression levels (RPKM values averaged over all individuals). For each codon, we compared its RSCU in the first (least expressed) and last (most expressed) class using a Mann-Whitney U test. Codons were annotated as preferred (resp. un-preferred) if their RSCU increased (resp. decreased) significantly with gene expression levels. All other codons were marked as non-significant. All synonymous SNPs for which an ancestral allele is unambiguously identified were annotated with regards to codon preference: mutations increasing codon preference (from un-preferred to either non-significant or preferred, or from non-significant to preferred) were annotated U→P while mutations decreasing codon preference (from preferred to either un-preferred or non-significant, or from non-significant to un-preferred) were annotated P→U. Mutations not affecting preference were considered as neutral with respect to SCU.

### substitutions

Using the three species alignments (Focal + two outgroups), we also counted and polarized substitutions specific to the focal species lineage. Divergent sites were determined as sites that were fixed in the focal population and different from both outgroups. Only sites identical in both outgroups were considered. As described above for SNPs, substitutions were classified as W→S, S→W or neutral, and U→P, P→U and neutral.

### Modified MK-test, neutrality and direction of selection indices

We performed a modified McDonald-Kreitman (MK) test [46], comparing W→S to S→W polymorphic and divergent sites on one hand (gBGC set) and U→P to P→U polymorphic and divergent sites on the other (SCU set). The underlying theory is detailed in Text S1. For each category, the total number of synonymous polymorphic and divergent sites was computed following the criteria detailed above. We performed a Chi-squared test for each set. Significant tests indicate that sequences do not evolve only under mutation pressure: selection and/or gBGC must be at work. Furthermore, we computed for each set a neutrality [47] and a direction of selection [48] indices as follows:

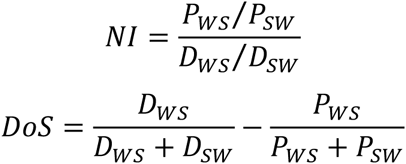

Where *P*_*WS*_ and *P*_*WS*_ are the number of W→S and S→W SNPs and *D*_*WS*_ and D_*WS*_ the number of W→S and S→W substitutions respectively. Assuming constant mutational bias, NI values lower than 1 or positive DoS values indicate SCU and/or gBGC of similar or stronger intensity at the divergence than at the polymorphism level. Respectively, NI values higher than 1, or negative DoS values indicate stronger selection and/or gBGC at the polymorphism than at the divergence level (see Sup. Text S1).

Because these statistics rely on polarized polymorphisms and substitutions, they are potentially sensitive to polarization errors, which could lead to spurious signature of selection/gBGC [33,40]. Importantly, we showed in Text S1 that the sign of both statistics is insensitive to polarization errors (as far as they are lower than 50%) and that polarization errors decrease the magnitude of the statistics, which makes our tests conservative to polarization errors.

### Estimation of gBGC and SCU

To estimate gBGC and SCU we extended the method of Glémin et al. [33] as detailed in Text S2. The rationale of the approach is to fit population genetic models to the three derived SFS including fixed mutations (W→S or U→P, S→W or P→U, and neutral). Parameters estimated are ancestral (*B*_0_ or *S*_0_) and recent (*B*_1_ or *S*_1_) gBGC or selection, mutational bias (λ), as well as other parameters (see Text S2 for details). We ran a series of nested models where *B*_0_ and *B*_1_ (or *S*_0_ and *S*_1_) are either fixed to zero or freely estimated, plus one model where they are set to be equal. Models were compared by the appropriate likelihood ratio tests (LRT). To jointly estimate gBGC and selection, we also extended the model by fitting nine SFS corresponding to the combination of the three basic SFS (e.g. W→S and P→U see Table S2.1 in Text S2 for the complete list). We tested all combinations of models where each parameter can be either null or freely estimated, so from the null neutral model, *B*_0_ = *B*_1_ = *S*_0_ = *S*_1_ = 0, to the model with the four parameters being freely estimated. As all models are not nested, we then chose the best model using the Akaike Information Criterium (AIC). When AICs were very close we chose the model with the lowest number of free parameters.

## Author contributions

### Conceived and designed the experiment

Sylvain Glémin and Jacques David

### Provided plant and molecular (RNA) resources

Roberto Bacilieri, Angélique Berger, Guillaume Besnard, Céline Cardi, Jacques David, Fabien De Bellis, Olivier Fouet, Bouchaib Khadari, Claire Lanaud, Thierry Leroy, David Pot, Christopher Sauvage, Nora Scarcelli, James Tregear, Yves Vigouroux, Nabila Yahiaoui

### Biomolecular work

Laure Sauné, Morgane Ardisson, Sylvain Santoni

### Processed data

Gautier Sarah, Yan Holtz, Felix Homa, Stephanie Pointet, Sandy Contreras, Benoit Nabholz, Roberto Bacilieri, Cyril Jourda, François Sabot, Manuel Ruiz

### Developed statistical models

Sylvain Glémin

### Performed analysis

Yves Clément and Sylvain Glémin

### Wrote the article

Yves Clément and Sylvain Glémin

### Administrated the project

Jean-Louis Pham and Jean-Pierre Labouisse

## Acknowledgements

We thank Nicolas Galtier for numerous discussions and for sharing scripts during the course of the project, Aurélien Bernard for help with bioinformatics, Carina Mugal and Laurent Duret for helpful comments on the manuscript. We thank the following colleagues and institutions for providing plant material: Dr Hyacinthe Legnate and the CNRA (Ivory Coast), Pr Adrien Kalondji and the University of Kinshasa (DRC), Bernard Perthuis (CIRAD French-Guiana) for Coffee-tree, Michel Boccara and the Cocoa Research Center (Trinidad) for Cocoa, Pierre-Oliver Cheptou and the staff of the CEFE experimental station CEFE-CNRS (Montpellier, France) for *Phyllirea* species, the Domaine de Vassal grapevine seed bank (INRA, Marseillan-Plage, France) for *Vitis* accessions, Frédérique Aberlenc for palm tree (Montpellier, France), CRB Plantes Tropicales Guadeloupe and CARBAP Cameroun for banana.

